# Network topology reveals distinct forms of developmental leverage in the Drosophila wing

**DOI:** 10.64898/2026.09.28.755080

**Authors:** Lanie Gardner, Dariann Adams, Kenneth Z. McKenna

## Abstract

Developmental gene regulatory networks reliably transform positional information into complex multicellular form, yet the organizational principles linking network architecture to developmental mechanism remain poorly understood. Here, we analyzed the *Drosophila melanogaster* wing developmental network to determine whether network topology reflects the distribution of developmental leverage during organogenesis. Integration of curated wing-development genes with high-confidence STRING interactions revealed five Hierarchical Layers of Developmental Control (HLDCs) associated with distinct topological and developmental roles. Organizer Centers, Signaling Scaffolds, and Pattern Implementers formed a forward-specification axis in which connectivity progressively contracted as positional information was transformed into increasingly localized developmental programs. Interface Coordinators departed from this hierarchy through disproportionate brokerage, whereas Local Modulators retained connectivity despite localized developmental scope. We propose that these complementary signatures reflect two regulatory architectures: 1) hierarchical information propagation that generates developmental identity and 2) distributed homeostatic regulation that coordinates and refines developmental outputs. Within Character Identity Modules (ChiMOs), this architecture links conserved patterning systems, Hox-defined contexts, and organ-specific kernels to reproducible morphology, providing a mechanistic hypothesis for developmental canalization and experimentally testable predictions.

**Summary Statement:** Developmental networks organize both the flow and preservation of information, revealing how distinct regulatory roles work together to generate precise, reproducible organ form.

## INTRODUCTION

The defining challenge of multicellular development is not simply to regulate genes, but to reliably transform developmental information into morphology. During organogenesis, cells must continuously determine where they are, what they should become, and how their behavior should remain coordinated with neighboring tissues despite ongoing growth, differentiation, stochastic molecular fluctuations, and environmental perturbation. Remarkably, these processes generate organs with highly reproducible size, shape, and organization across individuals. Understanding how developmental systems achieve this reproducibility remains one of the central challenges of developmental biology (Waddington, 1942; Félix and Barkoulas, 2015).

Gene regulatory networks (GRNs) have become the dominant framework for describing the molecular basis of development because they capture the regulatory interactions underlying cell specification and tissue patterning (Levine and Davidson, 2005; Davidson, 2010b). Decades of work have identified the signaling pathways, transcription factors, and regulatory circuits responsible for wing development in *Drosophila melanogaster*, establishing one of the best-characterized developmental systems in biology(Campbell and Tomlinson, 1998; Lecuit and Cohen, 1998; Lin and Perrimon, 1999; Tanimoto *et al*., 2000; Barrio and de Celis, 2004; Ruiz-Losada *et al*., 2018). Yet despite this detailed molecular knowledge, an important conceptual question remains unresolved. Although we understand many of the individual components of developmental networks, we know far less about how these components are organized into a coherent developmental process capable of reliably transforming positional information into morphology.

One limitation is that developmental genes are often studied in isolation or within individual signaling pathways(Gawne, McKenna and Nijhout, 2018; McKenna, Gawne and Nijhout, 2022). Perturbation experiments identify genes required for specific developmental events (Tanimoto *et al*., 2000), while network analyses quantify properties such as connectivity, centrality, or modularity (Albert and Barabasi, 2002; Rives and Galitski, 2003; Albert, 2007; Hintze and Adami, 2008). These approaches have been enormously successful in identifying developmental regulators, yet they rarely address what those network properties mean biologically. Highly connected genes are commonly described as “hubs,” but this designation is fundamentally descriptive rather than mechanistic. Connectivity itself does not explain why some genes influence the entire organ whereas others affect only highly localized aspects of morphology. Such differences in effect imply some form of network hierarchy.

Hierarchy in developmental GRNs has been defined in several related but non-equivalent ways. Classical developmental GRN models emphasize the causal ordering of regulatory states, in which upstream regulatory events establish conditions upon which progressively downstream developmental decisions depend (Davidson, 2010a; Peter and Davidson, 2017). In this sense, hierarchy describes what might be termed causal depth or the position of a regulatory interaction within the sequential transformation of developmental information. Other approaches have inferred hierarchy from the directed architecture, regulatory influence, or temporal relationships among regulators (Yu and Gerstein, 2006; Albert, 2007; Bhardwaj, Kim and Gerstein, 2010; Wu *et al*., 2021). These approaches have established hierarchy as a fundamental property of regulatory organization, but hierarchical position need not be equivalent to connectivity, information-flow control, or developmental influence.

Here, we consider a complementary dimension of developmental hierarchy - developmental leverage. We define leverage as the extent to which a component occupies a network position capable of establishing, transmitting, coordinating, implementing, or locally modifying developmental information during character formation. This distinction is important because genes at similar causal depths may occupy very different topological positions, whereas genes with comparable connectivity may perform fundamentally different developmental operations. We therefore treat degree, betweenness, and brokerage not as interchangeable measures of “importance,” but as partially independent properties that together describe how developmental influence is distributed through a character-generating network. Because the interaction network analyzed here is not a directed regulatory graph, our analysis does not attempt to reconstruct causal ordering among regulatory interactions. Instead, it asks whether topological position captures a complementary property—the distribution of developmental leverage across the network.

Here we test whether network topology captures this complementary dimension of developmental organization. We combined a manually curated set of canonical *Drosophila* wing developmental genes (Grimm and Pflugfelder, 1996; Campbell and Tomlinson, 1998; Lin and Perrimon, 1999; Ruiz-Losada *et al*., 2018) with high-confidence protein interaction data to quantify network connectivity. Rather than assigning hierarchy solely from upstream-to-downstream regulatory order, we quantified complementary properties of network position, including degree centrality, betweenness centrality, and residual brokerage (Yu *et al*., 2007; Koschützki and Schreiber, 2008; Pavlopoulos *et al*., 2011; Yaveroğlu *et al*., 2014). Multivariate analysis of these properties revealed recurrent topological regimes that showed substantial correspondence with established developmental functions. We interpret these regimes as five Hierarchical Layers of Developmental Control (HLDCs), representing the progressive establishment, transmission, coordination, implementation, and local refinement of developmental information.

To interpret these findings, we integrate the HLDC framework with the Character Identity Module (ChiMO) model (McKenna, 2021), which proposes that organ identity emerges through the interaction of conserved patterning systems, Hox-mediated developmental context, and organ-specific developmental kernels. Within this framework, the connectivity hierarchy identifies the information-processing architecture through which these regulatory systems interact during organogenesis. We propose that three hierarchical layers propagate developmental information from positional specification to regional cell identity, whereas two complementary layers maintain developmental integration and homeostasis throughout morphogenesis. This framework provides a mechanistic explanation for how developmental networks reliably generate reproducible morphology and suggests that canalization emerges from hierarchical information processing rather than from any single buffering mechanism.

## RESULTS

### Connectivity hierarchy reveals hierarchical layers of developmental control

Development proceeds through the progressive transformation of positional information into increasingly specialized cellular behaviors that ultimately generate adult morphology. Although gene regulatory networks provide a framework for describing the molecular interactions underlying this process, developmental genes are commonly considered either as components of individual signaling pathways or within a single regulatory network. Such approaches identify developmental interactions but do not necessarily reveal whether differences in network position correspond to distinct forms of developmental control. We therefore asked whether the interaction architecture of genes involved in *Drosophila* wing development contains a topological hierarchy corresponding to distinct mechanistic operations during organogenesis.

To address this question, we assembled a curated set of genes with experimentally established roles in patterning, growth, compartment specification, signaling, and differentiation of the *Drosophila* wing imaginal disc and mapped these genes onto the broader STRING interaction network. Rather than relying on a single measure of network position, we quantified complementary aspects of network topology. Degree centrality measured direct connectivity, betweenness centrality measured shortest-path positioning within the network, and residual brokerage identified genes whose betweenness was greater or lower than expected from their degree alone. Degree and betweenness were additionally integrated into a transformed connectivity hierarchy score representing relative position along the major core-to-periphery axis of the network.

### Wing developmental genes occupy a pronounced core-to-periphery connectivity hierarchy

Visualization of the wing developmental genes according to their connectivity hierarchy score revealed a pronounced core-to-periphery organization (Fig. 1A). A densely interconnected network core contained many canonical components of developmental signaling systems, including *dpp*, *hh*, *wg*, *N*, *arm*, *dsh*, *fz*, *smo* and *ci*, whereas progressively lower-scoring genes occupied increasingly peripheral network positions. Thus, genes involved in wing development were not distributed equivalently across the interaction network but instead occupied markedly different positions along a continuous connectivity hierarchy.

**Fig. 1.**
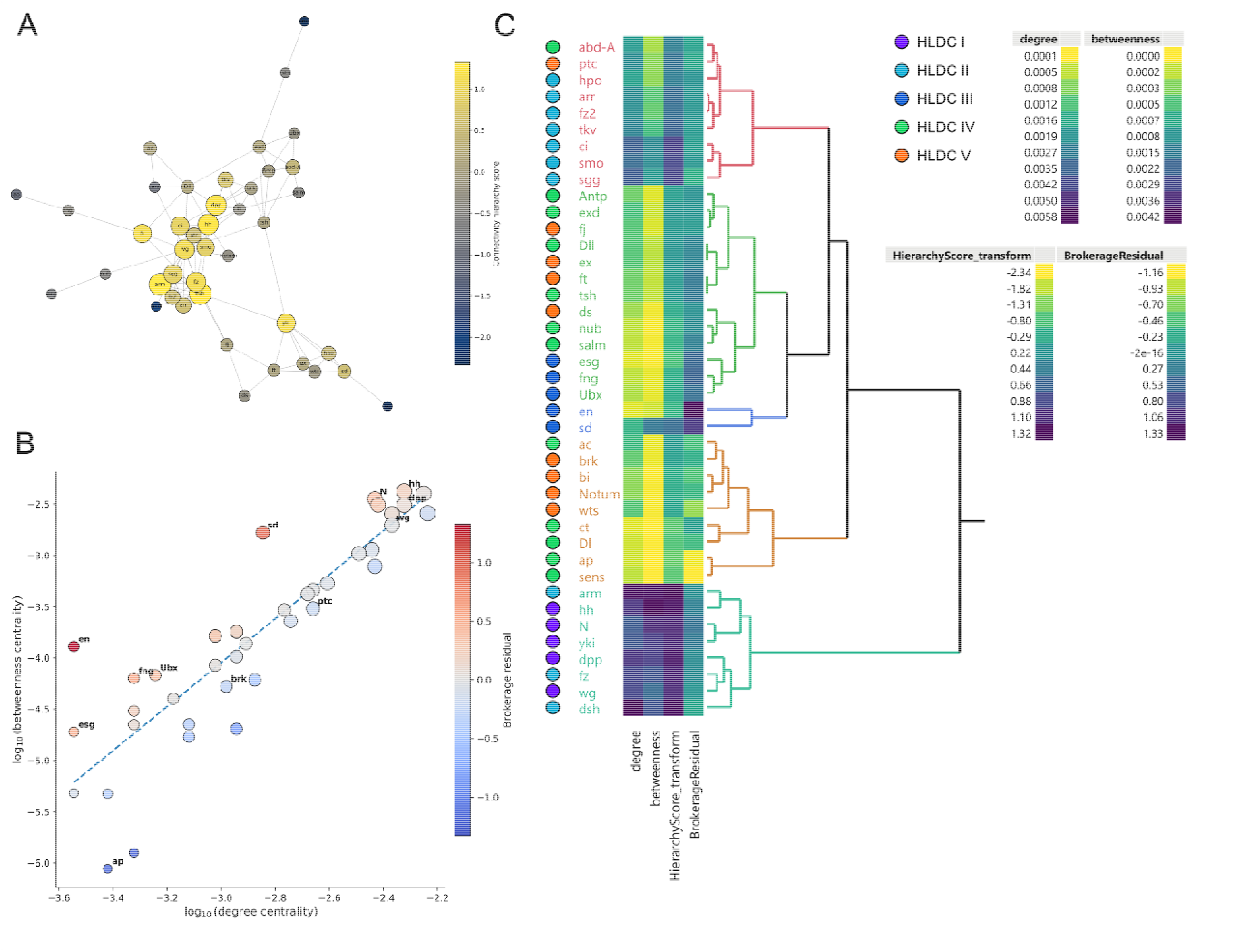
Connectivity architecture reveals hierarchical organization of the Drosophila wing developmental network. **(A)** Network representation of the curated wing developmental interaction network. Node size reflects degree centrality and node color represents transformed connectivity hierarchy score, revealing a densely connected developmental core surrounded by progressively more peripheral regulators. **(B)** Relationship between degree and betweenness centrality. The dashed line represents the expected degree–betweenness relationship, while point color indicates brokerage residual. Genes above the relationship exhibit greater betweenness than expected from their connectivity, identifying candidate regulatory brokers, including *en*, *sd*, *fng*, *Ubx*, and *esg*. **(C)** Hierarchical clustering of degree, betweenness, transformed hierarchy score, and brokerage residual reveals recurrent multivariate topological regimes. Colored circles denote independently assigned Hierarchical Layers of Developmental Control (HLDCs): HLDC I, Organizer Centers; HLDC II, Signaling Scaffolds; HLDC III, Interface Coordinators; HLDC IV, Pattern Implementers; and HLDC V, Local Modulators. Correspondence between clustering and developmental annotation indicates that network position is associated with distinct developmental operations, while overlap between classes demonstrates that HLDC identity reflects the integration of network architecture with developmental mechanism.

This organization was not attributable solely to the number of interactions associated with each gene. Degree and betweenness centrality were strongly related overall, but individual genes deviated substantially from the expected relationship between these metrics (Fig. 1B). Residuals from the log-degree–log-betweenness relationship therefore revealed a second dimension of network organization that was partially independent of overall connectivity. Several genes, including *en*, *fng*, *Ubx*, *esg* and *sd*, exhibited greater betweenness than expected from their degree and consequently high positive brokerage residuals. In contrast, other genes occupied positions below the expected degree–betweenness relationship. These results distinguish broadly connected nodes from nodes whose topological importance derives disproportionately from their position between different regions of the network.

This distinction was particularly important because genes with elevated residual brokerage were not necessarily those occupying the highest positions in the core-to-periphery hierarchy. For example, canonical signaling components were concentrated toward the highly connected region of the degree–betweenness relationship, whereas several genes subsequently classified as Interface Coordinators exhibited pronounced positive deviations despite more intermediate or peripheral connectivity (Fig. 1B). Network hierarchy therefore contained at least two distinguishable topological properties: overall connectivity position and disproportionate brokerage between network regions.

### Multivariate topology reveals recurrent network regimes corresponding to developmental function

We next asked whether these complementary topological properties jointly partitioned wing developmental genes into recurrent regions of network space. Hierarchical clustering of standardized degree centrality, betweenness centrality, transformed hierarchy score, and brokerage residual resolved a structured multivariate topology rather than a simple linear ordering of genes (Fig. 1C). The major branches separated a highly connected core from progressively less connected regions of the network, while residual brokerage further distinguished genes occupying unusual bridging positions within otherwise similar connectivity ranges.

Importantly, the clustering was performed on network metrics independently of the developmental annotations shown alongside the dendrogram. HLDC assignments were subsequently made by integrating topological position with experimentally established developmental function; the colored HLDC symbols in Fig. 1C therefore provide a visual comparison between unsupervised multivariate topology and independently curated biological interpretation rather than defining the clusters themselves. This comparison revealed substantial correspondence between recurrent topological regimes and developmental mechanism. Organizer Centers and Signaling Scaffolds were concentrated within the highly connected network core (Fig. 1A, Fig. S2), whereas Interface Coordinators were distinguished particularly by disproportionate brokerage (Fig. 1C, Fig. S3). Pattern Implementers extended furthest into the network periphery (Fig. 1A, Fig. S2), whereas Local Modulators occupied overlapping but generally more intermediate positions (Fig. 1A, Fig. S2), consistent with their continued participation in the signaling and regulatory processes whose outputs they refine. Correspondence was strongest among HLDCs I–III, whereas the lower-connectivity portion of the network showed greater topological overlap between HLDC IV and HLDC V.

Together, these analyses indicate that the interaction architecture of wing developmental genes cannot be described by connectivity alone. Instead, a broad core-to-periphery hierarchy is superimposed with a second brokerage dimension that identifies nodes occupying disproportionate bridging positions. The correspondence of these topological regimes with established developmental functions led us to propose five Hierarchical Layers of Developmental Control (HLDCs), defined as mechanistic developmental classes associated with characteristic regions of network topology.

At the broadest level, HLDC I (Organizer Centers) contains developmental organizers associated with establishment of tissue-wide positional information. HLDC II (Signaling Scaffolds) comprises components that receive, transmit, and interpret organizer-derived signals. HLDC III (Interface Coordinators) is distinguished by genes occupying disproportionate brokerage positions and functioning at interfaces among developmental programs. HLDC IV (Pattern Implementers) contains regional regulators that execute localized developmental programs and specify anatomical identities. HLDC V (Local Modulators) contains genes associated with localized regulation, refinement, and stabilization of developmental outputs. These categories should therefore not be interpreted as five discrete statistical clusters. Rather, they represent mechanistic classes associated with recurrent positions within a continuous, multidimensional network architecture (Fig. 1).

This organization suggests that connectivity hierarchy captures a property of developmental organization beyond simple pathway membership. High-connectivity positions are preferentially occupied by components associated with broad developmental signaling, whereas increasingly localized developmental functions occur toward the network periphery. Brokerage identifies an additional mode of network position in which genes may exert coordinative influence without correspondingly high overall connectivity. We therefore interpret these topological differences as reflecting distinct forms of developmental leverage as differences in the scale and manner by which components participate in the transformation of developmental information into morphology.

### The inferred connectivity hierarchy was robust to network-construction threshold

The relative positioning of wing developmental genes within the network was highly stable across STRING confidence thresholds (Table S1, Fig. S1). Pairwise Spearman correlations for the transformed hierarchy score ranged from *p=*0.860to 0.943across combined-score thresholds of 600–900 (all *p<*3.4 10^-12^). Degree centrality was similarly stable (*p=*0.895–0.969), while betweenness centrality showed somewhat greater but still substantial threshold sensitivity (*p=*0.754–0.890). Brokerage residuals were less stable (*p=*0.282–0.697), consistent with the greater sensitivity of shortest-path relationships to removal of lower-confidence interactions. Thus, the broad core-to-periphery connectivity hierarchy was robust to the STRING confidence threshold, whereas finer-scale estimates of brokerage were more threshold dependent.

### Summary of the connectivity hierarchy

Taken together, these results reveal two distinct forms of organization within the wing developmental network. A forward-specification axis comprising HLDCs I, II, and IV exhibits a pronounced connectivity hierarchy, with network position contracting from highly connected signaling components toward increasingly peripheral pattern implementers. In contrast, HLDCs III and V depart from this simple core-to-periphery progression in distinct ways. Interface Coordinators exhibit disproportionately high brokerage relative to their overall connectivity, whereas Local Modulators retain greater connectivity than expected from their localized developmental scope. Thus, the classes proposed to mediate developmental integration and homeostasis are distinguished not simply by their position along the connectivity hierarchy, but by characteristic departures from it. These complementary topological signatures suggest that developmental network architecture contains both a hierarchical axis associated with forward information propagation and a second regulatory architecture associated with the coordination and refinement of developmental outputs (Fig. 2, Fig. 3).

**Figure 2.**
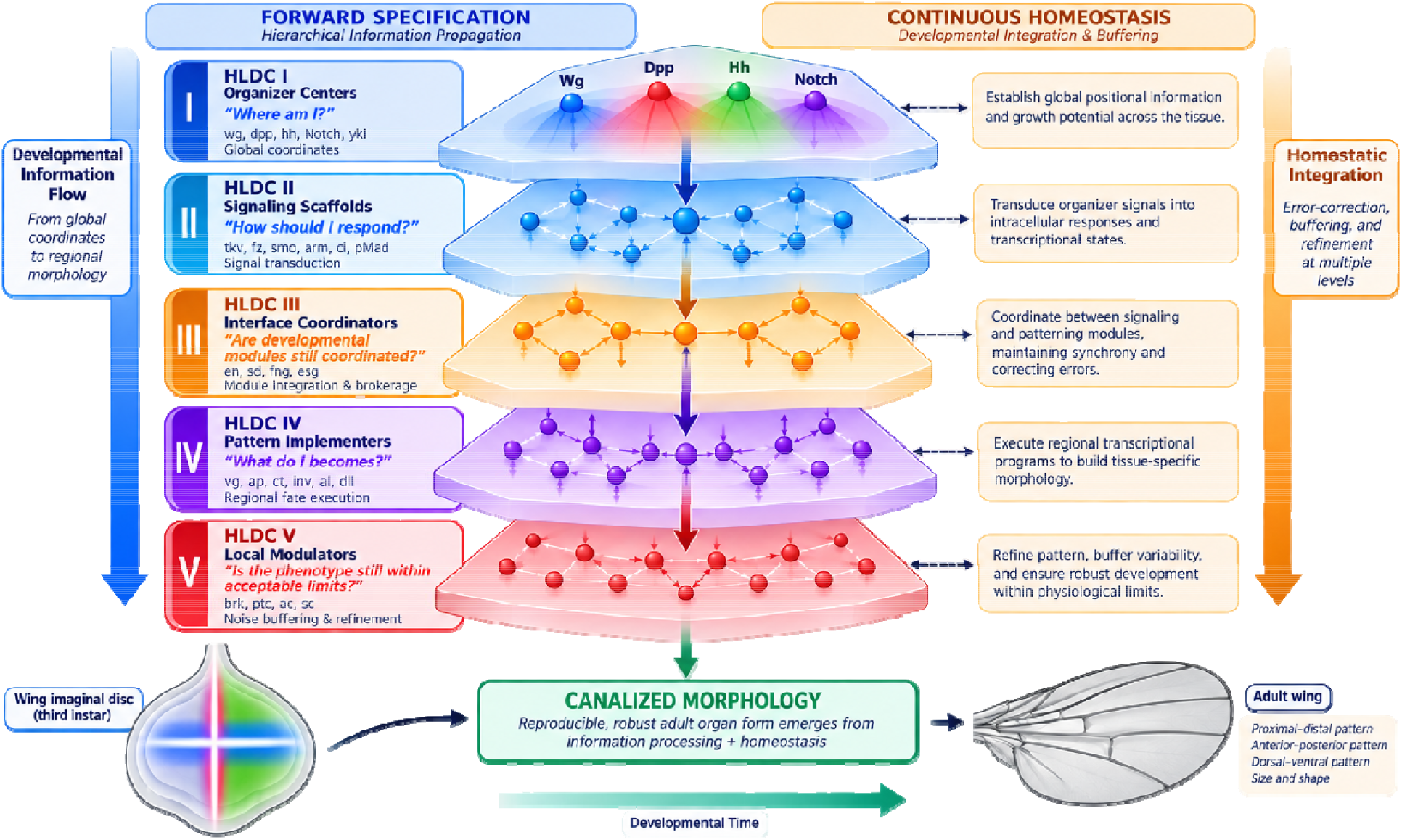
Dual regulatory architecture of the *Drosophila* wing developmental network. Organogenesis relies on two complementary, concurrently operating regulatory streams that converge to generate a reproducible, canalized organ form. (Left) The Forward Specification Axis drives hierarchical information propagation across three sequential layers: HLDC I (Organizer Centers) establishes tissue-wide spatial coordinates (“Where am I?”); HLDC II (Signaling Scaffolds) transduces extracellular signals into intracellular regulatory states (“How should I respond?”); and HLDC IV (Pattern Implementers) executes regional transcriptional kernels to commit cells to specific anatomical structures (“What do I become?”). (Right) The Continuous Homeostasis Axis operates in parallel to preserve system-wide developmental coherence and fidelity throughout morphogenesis. HLDC III (Interface Coordinators) acts as a high-brokerage regulatory bridge to synchronize semi-autonomous developmental modules across compartment boundaries (“Are developmental modules still coordinated?”), while HLDC V (Local Modulators) applies local negative feedback loops and thresholding to buffer stochastic molecular noise (“Is the phenotype still within acceptable limits?”). Dashed horizontal connectors illustrate lateral module synchronization and feedback refinement. Both regulatory axes converge upon Canalized Morphology, demonstrating that developmental robustness is an emergent property of simultaneous information specification and active homeostatic preservation. This figure was generated using ChatGPT (OpenAI) through iterative author-directed prompting beginning with an author generated schematic in PowerPoint. The final product was reviewed and revised by the author for scientific accuracy.

**Figure 3.**
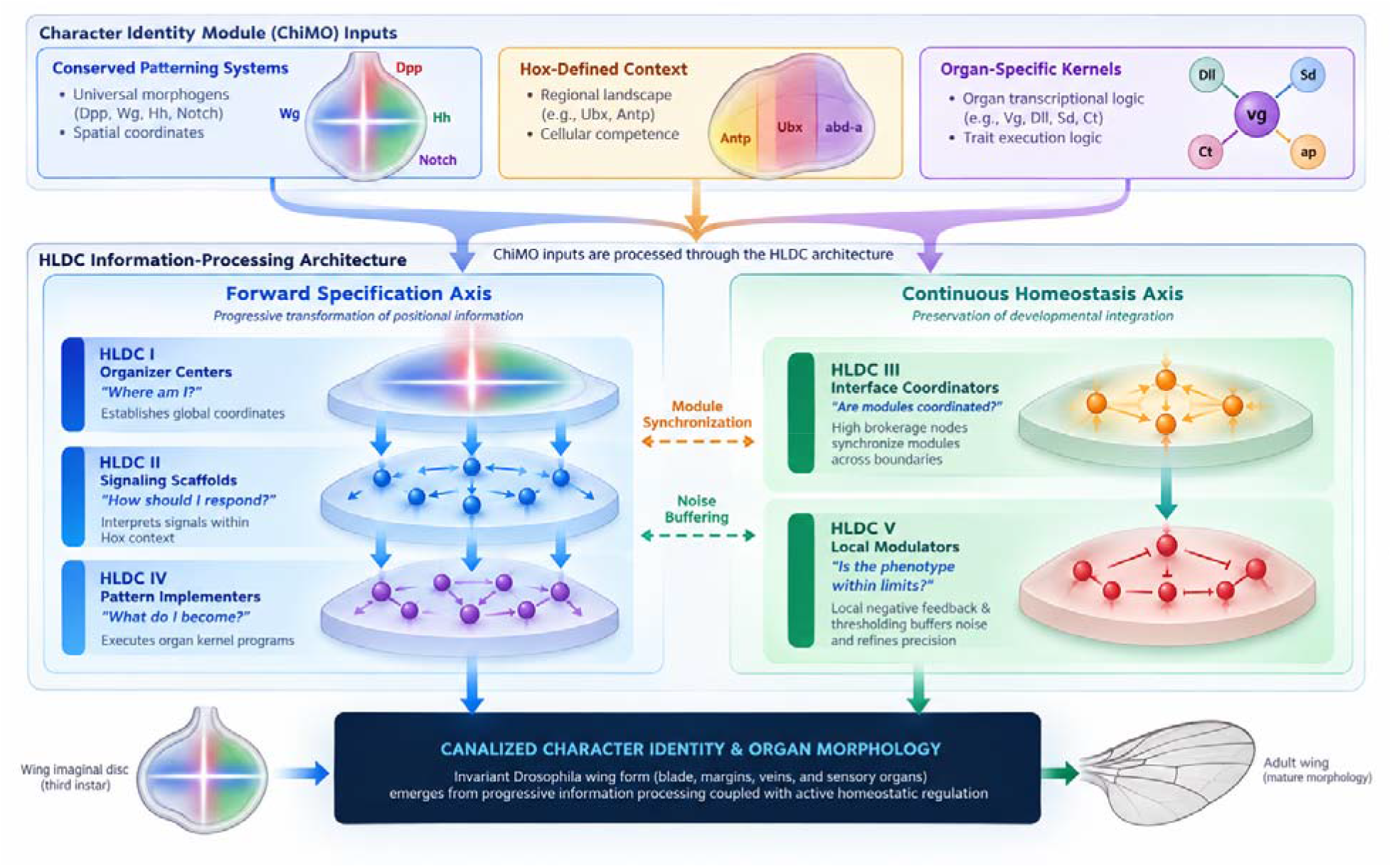
Integration of the Character Identity Module (ChiMO) model and the Hierarchical Layers of Developmental Control (HLDC) architecture. Organ identity emerges through the interaction of three primary ChiMO components: (i) conserved patterning systems that generate universal positional coordinates, (ii) Hox-defined developmental contexts that define regional regulatory landscapes, and (iii) organ-specific developmental kernels that execute localized trait-formation programs. The five HLDCs constitute the dynamic information-processing architecture through which these components are integrated during organogenesis. HLDC I (Organizer Centers) embeds universal patterning systems to establish spatial coordinates; HLDC II (Signaling Scaffolds) translates positional signals within Hox-defined regulatory contexts; HLDC III (Interface Coordinators) acts as a high-brokerage hub synchronizing communication among developing modules; HLDC IV (Pattern Implementers) expresses organ-specific developmental kernels to execute regional cell identities; and HLDC V (Local Modulators) buffers stochastic molecular noise to ensure phenotypic precision. Together, forward information propagation (HLDCs I, II, IV) and continuous homeostatic regulation (HLDCs III, V) explain how conserved signaling pathways reliably generate diverse, canalized organ identities. This figure was generated using ChatGPT (OpenAI) through iterative author-directed prompting beginning with an author generated schematic in PowerPoint. The final product was reviewed and revised by the author for scientific accuracy.

### HLDC I: Organizer Centers establish the developmental coordinate system

The first Hierarchical Layer of Developmental Control (HLDC I) comprises genes responsible for establishing the positional information that initiates wing development. Rather than specifying individual structures, these genes generate tissue-wide coordinate systems that enable every cell within the wing imaginal disc to determine its position relative to the developing organ. Because all subsequent developmental processes depend upon these positional cues, organizer genes occupy the highest positions within the connectivity hierarchy.

Genes assigned to HLDC I include the canonical developmental organizers wingless (wg), decapentaplegic (dpp), hedgehog (hh), Notch (N), and yorkie (yki) (Fig. 1C, Fig. 2). Together, these genes establish the primary developmental axes of the wing, regulate tissue growth, and generate the signaling environments that guide downstream developmental programs. Hedgehog produced in the posterior compartment induces dpp expression along the anterior-posterior boundary, while Wingless and Notch establish the dorsal-ventral organizer. Yorkie integrates growth control with these patterning mechanisms to coordinate tissue proliferation with positional specification (Fig. 2).

Consistent with their broad developmental roles, Organizer Centers exhibited the greatest degree and betweenness centrality within the developmental network, reflecting extensive regulatory interactions spanning multiple signaling pathways. This extensive connectivity is expected because organizer-derived positional information must be communicated throughout the developing organ before regional specification can occur.

The conceptual model presented in Figure 2 summarizes this developmental role. Organizer-derived morphogen gradients establish reproducible positional coordinates across the wing imaginal disc that are subsequently interpreted by downstream signaling pathways and regional transcriptional regulators. Thus, HLDC I represents the point at which developmental information enters the regulatory hierarchy, providing the positional framework upon which all subsequent patterning and morphogenesis depend.

### HLDC II: Signaling Scaffolds convert positional information into transcriptional responses

The second Hierarchical Layer of Developmental Control (HLDC II) comprises genes that receive organizer-derived signals and convert extracellular positional information into intracellular regulatory responses. Whereas HLDC I establishes the developmental coordinate system, HLDC II provides the molecular machinery that enables individual cells to detect, interpret, and respond to those positional cues.

Genes assigned to HLDC II include canonical components of the Wingless, Hedgehog, BMP, Hippo, and Notch signaling pathways, including receptors, cytoplasmic signaling proteins, and nuclear effectors. Representative members include frizzled (fz), frizzled2 (fz2), dishevelled (dsh), armadillo (arm), arrow (arr), smoothened (smo), thickveins (tkv), and cubitus interruptus (ci) (Fig. 1C, Fig. 2). Although these genes do not establish positional information themselves, they determine how organizer-derived signals are interpreted within responding cells.

Unlike Organizer Centers, whose activity is spatially restricted to discrete signaling domains, signaling scaffold components are broadly expressed throughout the developing wing epithelium. Their primary function is to receive extracellular morphogen signals and propagate them through intracellular signaling cascades, ultimately regulating developmental gene expression.

Signaling Scaffolds occupied relatively high positions within the connectivity hierarchy (Fig. S2), exhibiting extensive regulatory interactions across multiple developmental pathways. This network architecture is consistent with their central role in transmitting developmental signals from organizers to downstream transcriptional programs.

Collectively, these observations distinguish HLDC II from Organizer Centers despite their similarly high network connectivity. Organizer Centers establish positional information, whereas Signaling Scaffolds mediate its interpretation, representing a second, mechanistically distinct layer of developmental control.

### HLDC III: Interface Coordinators integrate developmental modules

Following the establishment and interpretation of positional information, successful organogenesis requires coordination among multiple developmental processes operating simultaneously within the developing wing. Growth, compartment identity, tissue boundaries, and regional patterning must remain synchronized despite being controlled by partially independent regulatory programs. These functions characterize the third Hierarchical Layer of Developmental Control (HLDC III).

Unlike Organizer Centers or Signaling Scaffolds, HLDC III genes were distinguished not by the greatest degree centrality but by disproportionately high brokerage relative to their overall connectivity (Fig. 1B, Fig. S3). Residual brokerage analysis identified these genes as regulatory bridges linking otherwise weakly connected regions of the developmental network, suggesting a role in coordinating communication between distinct developmental modules rather than broadcasting information throughout the entire network.

Representative Interface Coordinators included engrailed (en), scalloped (sd), fringe (fng), escargot (esg), and other regulators associated with compartment identity, boundary maintenance, and coordination among neighboring developmental programs (Fig. 2). Although these genes participate in fewer interactions than Organizer Centers, the interactions they maintain connect developmental processes that would otherwise remain only loosely integrated.

This network position is consistent with the modular organization of the wing imaginal disc, which comprises anterior and posterior compartments, dorsal and ventral territories, proliferative regions, and differentiating cell populations that must develop in a coordinated manner. As illustrated in Figure 2, Interface Coordinators occupy regulatory positions linking these partially autonomous modules, facilitating communication while preserving their functional independence.

The identification of HLDC III represents one of the principal biological insights emerging from the network analysis. Whereas degree centrality identified genes responsible for establishing and transmitting developmental information, brokerage identified a distinct class of regulators positioned to coordinate communication among semi-independent developmental modules. This finding suggests that connectivity hierarchy captures not only the propagation of developmental information but also the regulatory integration required to produce a coherent organ.

### HLDC IV: Pattern Implementers translate developmental information into regional cell identities

The fourth Hierarchical Layer of Developmental Control (HLDC IV) comprises genes that convert developmental information into stable patterns of cellular differentiation. Following establishment of positional information by Organizer Centers, its interpretation through Signaling Scaffolds, and coordination among developmental modules by Interface Coordinators, Pattern Implementers execute the localized developmental programs that generate the anatomical structures of the adult wing.

Genes assigned to HLDC IV function primarily as regional transcriptional regulators with spatially restricted expression domains. Representative members include vestigial (vg), apterous (ap), cut (ct), invected (inv), aristaless (al), and other transcription factors that specify distinct regions of the wing imaginal disc (Fig. 1C). Rather than generating or transmitting developmental information, these genes integrate upstream regulatory inputs to establish the identities of specific cell populations.

Unlike the broad developmental influence of Organizer Centers, Pattern Implementers act within discrete developmental territories (Fig. 2). Individual transcription factors regulate localized processes such as wing blade formation, dorsal-ventral identity, wing margin specification, sensory organ development, and vein differentiation. As illustrated in Figure 2, global positional cues established by upstream regulatory layers converge upon these regional regulators, which activate the gene expression programs appropriate for each developmental domain. In this way, continuous positional information is translated into discrete anatomical structures.

Consistent with their localized developmental roles, HLDC IV genes occupied lower positions within the connectivity hierarchy than Organizer Centers, Signaling Scaffolds, and Interface Coordinators. Their reduced network connectivity reflects the transition from organ-wide developmental regulation to the execution of regional differentiation programs, where extensive communication with the broader network is no longer required.

Together, these observations identify HLDC IV as the developmental layer in which positional information is committed to regional cell identity and morphology, transforming the regulatory information established by upstream layers into the distinct anatomical structures of the mature wing.

### HLDC V: Local Modulators refine developmental precision and robustness

The fifth Hierarchical Layer of Developmental Control (HLDC V) comprises genes that refine developmental outputs after positional information, regional identities, and tissue organization have already been established. Rather than initiating developmental programs, these genes regulate signaling thresholds, boundary precision, and local transcriptional responses to ensure accurate execution of morphogenesis.

Representative HLDC V genes include signaling antagonists, feedback regulators, threshold modifiers, and locally acting transcriptional regulators such as brinker (brk), patched (ptc), achaete (ac), and additional genes whose developmental effects are highly localized and context dependent (Fig. 1C). These genes function within established developmental environments to modulate signaling intensity, reinforce threshold responses, sharpen developmental boundaries, and stabilize cell fate decisions. As illustrated in Figure 2, Local Modulators act to refine rather than establishing developmental pattern.

Unlike the preceding layers, Local Modulators did not occupy a uniquely peripheral region of the connectivity hierarchy. Although HLDC V genes were generally displaced from the highly connected network core, their connectivity overlapped substantially with HLDC IV and in several cases exceeded that of terminal Pattern Implementers (Fig. 1C; Fig. S2). This distinction is consistent with their different developmental roles. Pattern Implementers can occupy highly peripheral positions because they execute increasingly regionalized transcriptional programs, whereas Local Modulators must remain coupled to the signaling systems whose outputs they refine. Consequently, modulation of a spatially restricted developmental process does not necessarily require minimal network connectivity.

This pattern is apparent among several representative HLDC V genes. *patched (ptc)* occupies a comparatively connected position because local regulation of Hedgehog signaling requires direct participation in the signaling architecture itself, whereas *brinker (brk)* occupies a more intermediate position consistent with its role in modulating transcriptional responses to Dpp signaling. *achaete (ac)* similarly lies outside the highly connected signaling core but is not among the most peripheral genes in the network, consistent with a regulatory role in localized pattern refinement rather than broad positional specification. Thus, the defining feature of HLDC V is not extreme network peripherality, but the combination of relatively restricted developmental scope with sufficient connectivity to modify, constrain, or refine locally operating developmental programs.

## DISCUSSION

### Connectivity hierarchy reveals the information-processing architecture of Character Identity Modules

One of the longstanding questions in evolutionary developmental biology is how a relatively conserved developmental toolkit repeatedly generates organs with remarkably different identities (Carroll, 2000, 2008). Morphogens such as Decapentaplegic (Dpp), Hedgehog (Hh), and Wingless (Wg) are deployed throughout animal development, yet these same signaling systems participate in the formation of wings, legs, antennae, eyes, and numerous other structures (Wolpert, 1969, 1971; Angelini and Kaufman, 2005). Likewise, Hox genes establish regional identity along the body axis but do not, by themselves, specify the detailed morphology of individual organs (Weatherbee and Carroll, 1999; Hughes and Kaufman, 2002; Pearson, Lemons and McGinnis, 2005; Lewis, 2007). Conversely, organ-specific developmental kernels provide the regulatory logic required for particular structures, yet they do not operate independently of broader positional information (Davidson, 2010a; McKenna, 2021). The developmental problem, therefore, is not simply one of identifying the genes involved in organ formation, but understanding how these conserved and organ-specific regulatory systems are integrated into a coherent developmental program.

To address this problem, we previously proposed the concept of the Character Identity Module (ChiMO) (McKenna, 2021), in which organ identity emerges through the interaction of three developmental components: (i) conserved patterning systems that establish positional coordinates, (ii) Hox genes that define the regional developmental landscape within which those coordinates are interpreted, and (iii) organ-specific developmental kernels that translate positional information into distinct cellular identities and morphological traits. Within this framework, morphogens provide a common positional reference system shared across organs, Hox genes modify the regulatory context in which those positional signals are interpreted, and developmental kernels determine the organ-specific transcriptional programs that ultimately generate unique morphologies. Character identity is therefore not encoded by any single regulatory network, but emerges from the coordinated interaction among these three developmental systems.

The present analysis extends this conceptual framework by proposing that the five Hierarchical Layers of Developmental Control (HLDCs) constitute the information-processing architecture through which ChiMOs operate. Rather than representing independent functional categories, the HLDCs describe successive transformations of developmental information during organogenesis. Organizer Centers establish positional information, Signaling Scaffolds determine how cells interpret those positional cues, Interface Coordinators integrate communication among developing cellular populations, Pattern Implementers execute organ-specific differentiation programs, and Local Modulators refine developmental outputs through local feedback and phenotypic stabilization. Collectively, these layers transform inherited developmental information into stable morphology.

The concept of developmental leverage provides a way to interpret this architecture. Whereas causal depth describes where a regulatory interaction occurs within an upstream-to-downstream sequence (Yu and Gerstein, 2006; Bhardwaj, Kim and Gerstein, 2010; Davidson, 2010a; Peter and Davidson, 2017), developmental leverage describes the scale and character of developmental influence associated with a component’s position in the network. Organizer Centers possess broad leverage because their activity contributes to positional information used across large regions of the developing organ. Signaling Scaffolds transmit that influence into cellular regulatory states, while Pattern Implementers increasingly restrict it to regional developmental programs. Interface Coordinators represent a different form of leverage. Their importance derives not primarily from broad connectivity, but from their disproportionate position between otherwise semi-independent developmental modules. Local Modulators occupy the most spatially restricted level, where developmental influence is concentrated on the precision and stability of local outputs. The HLDCs therefore describe not simply a sequence of regulatory events, but a progressive redistribution of developmental leverage as positional information is transformed into morphology.

Viewed in this way, ChiMOs are not static collections of interacting genes but dynamic information-processing systems. Positional information established by conserved morphogens is progressively interpreted, integrated, and refined as development proceeds (Wolpert, 1969; Tkačik and Gregor, 2021). At each hierarchical layer, developmental information becomes increasingly specific. Broad spatial coordinates give rise to regional developmental environments, those environments specify distinct cellular identities, and coordinated cellular behaviors ultimately generate organ morphology. Connectivity hierarchy therefore reflects more than patterns of molecular interaction. It reveals differences in developmental leverage associated with the regulatory transformations through which developmental information is progressively converted into form.

Importantly, this framework also resolves an apparent paradox in developmental biology. Conserved signaling pathways can participate in the formation of many different organs because they do not directly encode organ identity. Instead, they establish a common coordinate system that is interpreted differently depending on the surrounding Hox environment and the organ-specific developmental kernel (Wolpert, 1971; Davidson, 2010a; McKenna, 2021). The HLDC framework provides the mechanistic architecture through which these interactions occur. Rather than acting as isolated pathways, conserved patterning systems, Hox genes, and developmental kernels become components of a hierarchical developmental computation that progressively transforms positional information into character identity.

From this perspective, the primary function of a developmental gene regulatory network is not simply to regulate gene expression, but to regulate the progressive transformation of developmental information into morphology. Genes constitute the molecular substrate through which this transformation is accomplished, while connectivity hierarchy reflects the organization of the computational process itself. Organogenesis therefore emerges as the hierarchical processing of developmental information through an integrated regulatory architecture, providing a mechanistic foundation for both the stability of multicellular form.

### Forward developmental information propagation through Organizer Centers, Signaling Scaffolds, and Pattern Implementers

The HLDC framework suggests that organogenesis proceeds through a series of hierarchical regulatory transformations that progressively convert positional information into morphology (Wolpert, 1969; Peter and Davidson, 2017; Tkačik and Gregor, 2021). The first, second, and fourth hierarchical layers collectively function as the forward information-processing architecture of development (Fig. 3). Rather than representing independent regulatory processes, these layers perform successive developmental computations in which broad positional information is transformed into increasingly specific cellular identities and morphological outcomes.

### HLDC I: Organizer Centers establish developmental coordinate systems

Development begins by establishing spatial reference frames that allow cells to determine their position within the developing organ (Wolpert, 1969; Tkačik and Gregor, 2021). Organizer Centers occupy the highest level of the developmental hierarchy because they generate the coordinate systems upon which all subsequent developmental decisions depend. Classical organizer genes such as dpp, hh, and wg establish morphogen gradients that partition developing tissues into distinct positional domains and provide cells with information about their relative location within the organ (Lecuit and Cohen, 1998; Tanimoto *et al*., 2000; Harmansa *et al*., 2017; Ruiz-Losada *et al*., 2018).

Organizer signals generally act upstream of the transcriptional programs that specify final cell identities and morphological structures (Wolpert, 1969; Ruiz-Losada *et al*., 2018; Tkačik and Gregor, 2021). Instead, they answer the fundamental developmental questions (Fig. 2): Where is a cell located? What positional information is available? These positional cues provide a common developmental framework that is reused throughout metazoan development. Consequently, the remarkable conservation of organizer pathways across organs and phyla reflects not conservation of morphology, but conservation of the positional coordinate systems required for multicellular pattern formation (Panganiban *et al*., 1997).

Within the ChiMO framework (Fig. 3), Organizer Centers therefore constitute the universal patterning systems that establish the spatial foundation upon which organ-specific developmental programs are subsequently constructed (McKenna, 2021).

### HLDC II: Signaling Scaffolds interpret positional information

Positional information alone is insufficient to generate morphology. Cells must also determine how they should respond to the developmental coordinates established by Organizer Centers. Signaling Scaffolds perform this second hierarchical transformation by regulating signal reception, intracellular transduction, transcriptional responsiveness, and developmental competence (Lecuit and Cohen, 1998; Lin and Perrimon, 1999; Tanimoto *et al*., 2000). Rather than generating positional information themselves, these genes determine how positional information is interpreted by individual cells.

Accordingly, Signaling Scaffolds answer the next set of developmental questions (Fig. 2): How should a cell respond to its position? Which signaling state should it adopt? The same morphogen concentration can therefore elicit different developmental outcomes depending upon the regulatory environment established by these signaling components (Wolpert, 1969). In this manner, positional information is transformed from a spatial coordinate into a specific cellular regulatory state.

This interpretation layer also provides a mechanistic bridge to the Hox component of the ChiMO framework (Fig. 3). Hox genes modify the regulatory landscape within which signaling pathways operate by altering cellular competence, transcriptional responsiveness, and signaling dynamics (Rezsohazy *et al*., 2015). Consequently, identical positional information can be interpreted differently in distinct Hox environments, allowing conserved morphogen systems to participate in the development of diverse morphological structures.

### HLDC IV: Pattern Implementers execute organ-specific developmental programs

Once positional information has been established and interpreted, development proceeds to the execution of organ-specific morphogenesis. Pattern Implementers occupy this fourth hierarchical layer by translating developmental states into stable cellular identities, differentiation programs, and tissue-specific behaviors. Genes within this class regulate the local developmental processes that directly generate morphology through changes in cell fate, polarity, proliferation, differentiation, and tissue architecture (Davidson, 2010a; Peter and Davidson, 2017).

These genes answer the final developmental questions of forward information propagation (Fig. 2): What does this cell become? How is morphology generated? At this stage, developmental information has progressed from abstract positional coordinates to concrete biological structures.

Within the ChiMO framework (Fig. 3), Pattern Implementers represent the level at which organ-specific developmental kernels are expressed. Developmental kernels do not generate positional information themselves; rather, they interpret positional information within a particular Hox-defined developmental context to produce unique morphological outcomes. For example, the limb developmental kernel centered on Distal-less (Dll) operates within positional axes established by broader patterning systems rather than independently generating those axes (Panganiban, Nagy and Carroll, 1994; Panganiban *et al*., 1997; Panganiban and Rubenstein, 2002; McKenna, 2021). Instead, it reads the positional information established by conserved morphogen systems and converts those developmental coordinates into the transcriptional programs that generate appendage identity. Thus, developmental kernels function as the organ-specific computational machinery that transforms conserved positional information into novel morphological characters.

### Forward developmental information propagation as hierarchical computation

Together, Organizer Centers, Signaling Scaffolds, and Pattern Implementers constitute the forward information-processing pathway of organogenesis. Development proceeds through successive regulatory transformations in which spatial coordinates are established, interpreted within a context-dependent regulatory environment, and ultimately translated into stable cellular identities and tissue morphology. Across these layers, developmental leverage progressively contracts. Broadly distributed positional information is transformed into increasingly restricted regulatory states and ultimately into local programs of cellular differentiation. This contraction does not imply decreasing developmental importance. Rather, it reflects a change in the spatial and regulatory scale at which developmental information acts.

Importantly, this hierarchy also provides a mechanistic explanation for the remarkable conservation of early developmental pathways alongside the extraordinary diversity of animal morphology (Panganiban, Nagy and Carroll, 1994; Panganiban *et al*., 1997; Carroll, 2008; McKenna, 2021). Organizer Centers and Signaling Scaffolds provide broadly conserved computational operations that establish and interpret positional information (Angelini and Kaufman, 2005), whereas Pattern Implementers—including organ-specific developmental kernels—translate these conserved developmental inputs into the unique cellular behaviors that define individual organs. In this view, morphological diversity emerges not because developmental systems use fundamentally different organizer pathways, but because conserved developmental information is interpreted and executed differently within distinct regulatory and evolutionary contexts (Panganiban *et al*., 1997; McKenna, 2021).

Forward information propagation alone, however, is insufficient to explain the remarkable reproducibility of development. As cellular populations diversify and developmental modules become increasingly specialized, additional regulatory mechanisms are required to coordinate their interactions and prevent errors from propagating through the developing organ.

### Developmental homeostasis emerges through Interface Coordinators and Local Modulators

Forward propagation of developmental information alone cannot account for the extraordinary reproducibility of multicellular development. As organogenesis progresses, increasingly specialized cell populations emerge, developmental modules become progressively more independent, and local interactions multiply throughout the developing organ. Without additional regulatory mechanisms, small stochastic fluctuations or local developmental errors would accumulate, propagating through the network and ultimately compromising organ morphology (Kerszberg, 2004; Félix and Barkoulas, 2015; Wong and Gilmour, 2020). Our analyses suggest that the third and fifth Hierarchical Layers of Developmental Control represent a complementary regulatory architecture whose primary role is not to specify new cellular identities, but to preserve the integrity of developmental trajectories as morphogenesis unfolds.

### HLDC III: Interface Coordinators synchronize independently developing modules

Interface Coordinators occupy a unique position within the developmental hierarchy because they regulate communication among developmental modules rather than specifying the identity of any single module itself. Genes within this class—including classical boundary organizers and selector genes such as engrailed (en) and fringe (fng)—maintain developmental coherence by coordinating interactions among neighboring cellular populations that are simultaneously undergoing distinct developmental programs (Tabata *et al*., 1995; Domínguez and Celis, 1998; Irvine, 1999; Rauskolb, Correia and Irvine, 1999; Irvine and Rauskolb, 2001; Joyner, Ortigão-Farias and Kornberg, 2024).

These genes answer a unique question among the HLDC levels (Fig. 2): Are developmental modules still coordinated? Importantly, these genes should not be viewed simply as regulators of compartment boundaries. Rather, compartment boundaries represent one manifestation of a more general developmental function - preserving communication among increasingly specialized developmental modules.

As development proceeds, distinct cellular populations begin executing independent regulatory programs while remaining physically and functionally integrated within the same organ (Irvine and Rauskolb, 2001; Joyner, Ortigão-Farias and Kornberg, 2024). Interface Coordinators help maintain synchronization among these parallel developmental trajectories by regulating the exchange of positional information, signaling activity, and developmental state across module interfaces.

The topology of Interface Coordinators suggests a qualitatively different form of developmental leverage. Unlike Organizer Centers, their influence does not arise primarily from high overall connectivity. Instead, their elevated brokerage indicates that they occupy positions through which otherwise partially separable regions of the developmental network remain connected (Yu *et al*., 2007; Bhardwaj, Kim and Gerstein, 2010). We refer to this as coordinative leverage or the capacity of a component to influence developmental coherence by regulating interactions among modules rather than by broadly distributing positional information.

From this perspective, we propose that Interface constitute a distributed system of developmental error correction (Fig. 2, Fig. 3). By regulating interactions among neighboring developmental modules, they may prevent local inconsistencies from propagating throughout the developing organ. Their importance therefore lies not in generating developmental information, but in maintaining the consistency and integration of developmental information as increasingly complex regulatory programs unfold.

Within the ChiMO framework (Fig. 3), this coordination is essential because organ identity emerges from the interaction of multiple regulatory systems operating simultaneously (McKenna, 2021). Conserved morphogen gradients, Hox-defined developmental contexts, and organ-specific developmental kernels cannot function independently. Their outputs must remain synchronized across space and time if a coherent morphological structure is to emerge. We propose that Interface Coordinators provide part of the regulatory architecture through which these independent developmental processes remain integrated into a single developmental program.

### HLDC V: Local Modulators refine developmental precision

Whereas Interface Coordinators regulate communication among developmental modules, Local Modulators operate at the level of individual cellular populations to ensure developmental precision. These genes regulate signaling thresholds, local feedback loops, boundary refinement, and quantitative aspects of cellular behavior that collectively determine the fidelity of the final phenotype (Kirkpatrick, Johnson and Laughon, 2001; Torroja, Gorfinkiel and Guerrero, 2004; Gallet and Therond, 2005; Schwank, Restrepo and Basler, 2008).

Rather than establishing developmental identities, Local Modulators fine-tune developmental outcomes by reducing local variability in cellular responses. Small fluctuations in signaling intensity, transcriptional activity, or cell behavior inevitably arise during development through stochastic molecular processes and environmental variation (Kerszberg, 2004; Félix and Barkoulas, 2015). We propose Local Modulators buffer such fluctuations before they can propagate into larger-scale morphological defects, thereby increasing developmental precision without fundamentally altering developmental identity.

These genes answer a more nuanced question among the HLDC levels (Fig. 2, Fig. 3): Is the phenotype still within acceptable limits? Rather than distributing information broadly across the developing organ, their influence is concentrated within particular signaling environments, cellular populations, or morphological domains. We propose that this localization allows developmental systems to adjust thresholds, sharpen boundaries, and stabilize outputs without requiring changes to the broader positional architecture from which those states were derived.

Consequently, Local Modulators may represent a local stage of developmental homeostasis, where broad developmental programs are refined into the precise, reproducible morphological features characteristic of a species.

### Developmental homeostasis as a hierarchical regulatory architecture

Together, Interface Coordinators and Local Modulators constitute a distributed homeostatic architecture that stabilizes developmental trajectories throughout organogenesis (Fig. 3). Unlike the forward information-processing layers, which progressively generate positional information, cellular identity, and morphology, these regulatory classes continuously integrate and refine developmental processes operating across multiple spatial and temporal scales. The HLDC framework therefore suggests that developmental robustness may emerge not as a passive consequence of network complexity, but as an active property of hierarchical regulatory organization (Siegal and Bergman, 2002; Félix and Barkoulas, 2015; Hallgrimsson *et al*., 2019).

This perspective provides a mechanistic hypothesis of canalization (Waddington, 1942; Siegal and Bergman, 2002; Hallgrimsson *et al*., 2019). Rather than requiring a single buffering mechanism, the HLDC framework predicts that canalization can emerge through coordinated regulatory feedback operating across multiple hierarchical levels. Interface Coordinators preserve coherence among independently developing modules, while Local Modulators suppress local developmental noise before it propagates through the regulatory hierarchy. Together, these complementary regulatory systems ensure that the forward propagation of developmental information consistently converges upon the species-typical phenotype despite stochastic molecular variation, environmental perturbation, and genetic diversity (Fig. 3).

Viewed in this manner, organogenesis is best understood as the inheritance and execution of a spatio-temporal homeostatic system. Development reliably establishes positional coordinates, specifies distinct cellular populations, regulates communication among those populations, and continuously stabilizes developmental trajectories through distributed positive and negative feedback (Siegal and Bergman, 2002; Félix and Barkoulas, 2015; Hallgrimsson *et al*., 2019). The remarkable reproducibility of multicellular form is therefore not an intrinsic property of individual developmental pathways, but an emergent consequence of hierarchical information processing coupled with hierarchical homeostatic regulation.

### Study Limitations

Several limitations of the present analysis should be considered when interpreting the HLDC framework. First, the network analyzed here is derived from high-confidence STRING interactions among a manually curated set of genes with established roles in *Drosophila* wing development. STRING integrates multiple classes of molecular association and does not constitute a directed, stage-specific regulatory network of the developing wing (Szklarczyk *et al*., 2025). Consequently, the topology analyzed here cannot reconstruct causal regulatory order, distinguish activating from inhibitory interactions, or capture the dynamic changes in network structure that occur across developmental time. Our measures should therefore be interpreted as properties of interaction topology rather than direct measurements of regulatory causality. The curated gene set also necessarily reflects the depth of existing experimental knowledge and may underrepresent poorly characterized components of wing development. Nevertheless, the major core-to-periphery organization was preserved across STRING confidence thresholds ranging from 600 to 900, indicating that the principal connectivity hierarchy is not dependent upon a single interaction-score cutoff.

Second, the developmental functions assigned to the HLDCs represent mechanistic interpretations of network position rather than experimentally validated properties of the classes themselves. Hierarchical clustering revealed recurrent topological regimes that showed substantial correspondence with established developmental functions, but the HLDCs should not be interpreted as synonymous with statistical clusters. In particular, the proposed roles of Interface Coordinators in developmental integration and Local Modulators in homeostatic stabilization generate predictions that require direct experimental testing. The framework predicts that standardized perturbation across hierarchical layers should reveal systematic differences in the spatial scope and character of developmental effects. Disruption of high-leverage components should propagate across broader regions of the developing organ, whereas perturbation of increasingly localized components should produce correspondingly restricted effects. It further predicts that disruption of Interface Coordinators or Local Modulators should disproportionately compromise developmental consistency, for example by increasing spatial discordance, phenotypic variance, or sensitivity to genetic and environmental perturbation. These predictions provide explicit experimental tests by which the proposed relationship between network topology, developmental leverage, and canalization can be evaluated.

## Conclusion

Development has traditionally been viewed as a process through which cells progressively acquire increasingly specialized identities. Our analyses suggest that this perspective captures only half of the developmental problem. Organogenesis requires not only the generation of developmental information but also the continuous preservation of that information as increasingly specialized cellular populations emerge. Developmental leverage provides a means of describing how these operations are distributed across the network. Broad positional influence is progressively transformed into increasingly localized developmental programs, while complementary coordinative and modulatory interactions preserve coherence as that transformation occurs. The HLDC framework therefore identifies two complementary regulatory architectures operating simultaneously throughout development - one that progressively specifies cellular identity through hierarchical information propagation and another that continuously integrates, refines, and stabilizes those developmental trajectories through hierarchical homeostatic regulation. Canalization therefore emerges not as a passive property of developmental systems but as the inevitable consequence of an architecture that continuously verifies and preserves developmental identity while it is being generated.

We propose that the fundamental organizational principle of multicellular development is not gene regulation alone, but the hierarchical generation and preservation of developmental information through an integrated homeostatic system.

## MATERIALS AND METHODS

### Curated *Drosophila melanogaster* wing developmental gene set

A curated set of canonical *Drosophila melanogaster* wing developmental genes was assembled from established literature describing wing pattern formation, growth regulation, compartment specification, selector gene activity, and tissue differentiation (Campbell and Tomlinson, 1998; Lecuit and Cohen, 1998; Lin and Perrimon, 1999; Tanimoto *et al*., 2000; Barrio and de Celis, 2004; Ruiz-Losada *et al*., 2018). The curated gene set included components of major developmental signaling pathways, transcriptional regulators, growth-control genes, and genes with experimentally characterized roles in wing morphogenesis.

Gene inclusion was based on published developmental evidence and was performed independently of the network metrics calculated in this study. Genes were selected based on their demonstrated involvement in establishing, interpreting, integrating, or refining developmental information within the wing imaginal disc. The initial curated dataset contained 52 genes, of which 45 were successfully mapped to STRING identifiers and included in network analyses. Gene identities and manually curated developmental annotations are provided in the accompanying dataset.

### STRING protein interaction network construction

To characterize the network position of wing developmental genes within a broader molecular interaction landscape, protein association data were downloaded from the STRING database for *Drosophila melanogaster* (Szklarczyk *et al*., 2025). The STRING interaction network was constructed using high-confidence interactions with a combined interaction score threshold of ≥700.

Because developmental genes function within a larger cellular network rather than as an isolated pathway, network topology was calculated using the full STRING interactome rather than a reduced wing-specific subnetwork. This approach allowed the analysis to capture the relative connectivity and network position of wing developmental genes within the broader molecular interaction architecture.

Gene symbols from the curated wing developmental dataset were mapped to STRING protein identifiers using the STRING protein information table linking preferred gene names to STRING protein IDs. Only genes with successful STRING mappings were included in network topology analyses. Additional curated wing genes lacking STRING mappings were retained for developmental classification when appropriate.

### Network-threshold sensitivity analysis

To determine whether inferred network position depended strongly on the STRING interaction-confidence threshold, the complete analysis was repeated using combined-score thresholds of 600, 700, 800, and 900. Threshold-specific datasets contained 45, 45, 44, and 39 mapped wing-development genes, respectively. Stability of network position was assessed using pairwise Spearman rank correlations between thresholds for degree centrality, betweenness centrality, transformed hierarchy score, and brokerage residual. Genes were matched by gene symbol between each pair of thresholds, and correlations were calculated using genes for which the corresponding metric was available at both thresholds. Because brokerage residual could only be calculated for genes with nonzero degree and betweenness, sample sizes for brokerage comparisons were smaller than those for the other metrics. All six pairwise combinations of the four STRING thresholds were evaluated.

### Network topology analysis

Network topology was quantified using degree centrality and betweenness centrality (Freeman, 1977; Koschützki and Schreiber, 2008; Pavlopoulos *et al*., 2011). Both measures were calculated on the complete STRING interaction network at the specified combined-score threshold, after which values corresponding to the curated wing developmental genes were extracted for subsequent analysis. Degree centrality was calculated using NetworkX (Hagberg, Swart and Schult, 2008) as the fraction of nodes in the network to which each node was directly connected. Because exact betweenness calculation across the full STRING network was computationally intensive, betweenness centrality was estimated using the NetworkX node-sampling algorithm with 1,000 sampled nodes (k = 1000) and a fixed random seed (seed = 42) to ensure reproducibility. Betweenness centrality estimates the proportion of shortest paths passing through a node and therefore identifies nodes occupying potential bridging positions between otherwise separated regions of the network.

Degree and betweenness centrality capture complementary properties of network position: degree quantifies direct connectivity, whereas betweenness quantifies shortest-path positioning within the broader network. These measures were subsequently integrated to characterize connectivity hierarchy and analyzed separately to identify nodes exhibiting disproportionate brokerage relative to their overall connectivity.

### Transformed connectivity hierarchy score

To characterize the combined contribution of direct connectivity and shortest-path position, we defined a composite connectivity hierarchy score integrating degree and betweenness centrality. Because both measures were strongly right-skewed and some genes exhibited zero betweenness, values were transformed according to

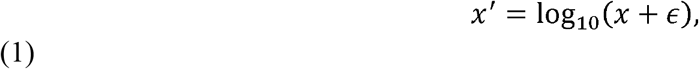

Where ℇ=10^-8^. Transformed degree and betweenness values were independently standardized as z-scores using their respective means and sample standard deviations. The transformed connectivity hierarchy score for each gene was then calculated as

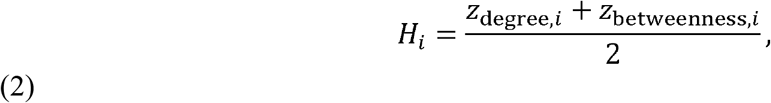

giving equal weight to transformed degree and betweenness centrality. This transformation retained genes with zero betweenness while reducing the influence of highly skewed centrality distributions. Genes were ranked according to the resulting hierarchy score and divided into five descriptive connectivity levels using quintile-based classification (pandas.qcut,*q*=5). These connectivity levels were used to summarize relative network position and were treated independently from subsequent mechanistic assignment to HLDC classes.

### Brokerage residual analysis identifies developmental integration nodes

Because genes with high connectivity and genes with high integrative capacity may represent distinct developmental roles (Yu *et al*., 2007; Bhardwaj, Kim and Gerstein, 2010), brokerage was analyzed independently from overall connectivity. We defined brokerage residual as the residual relationship between log-transformed degree and log-transformed betweenness.

To distinguish direct connectivity from disproportionate shortest-path positioning, brokerage was quantified from the relationship between degree and betweenness centrality. Genes with nonzero degree and betweenness values were retained and both measures were log10-transformed. An ordinary least-squares line was fitted in log-log space using numpy.polyfit, such that

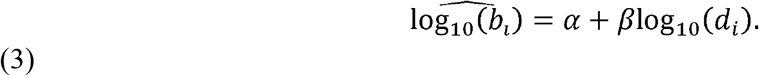

Brokerage residual was then defined as

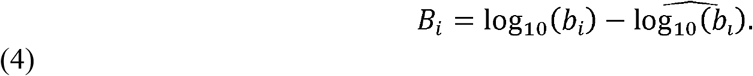

Positive residuals therefore identify nodes with greater betweenness than expected from their degree, whereas negative residuals identify nodes with lower betweenness than expected from their degree. Genes with zero betweenness were excluded from brokerage-residual calculations because their values could not be log-transformed, but were retained in analyses of connectivity hierarchy.

### Hierarchical clustering

To evaluate whether wing developmental genes occupied recurrent multivariate regions of network topology, hierarchical clustering was performed in JMP using degree centrality, betweenness centrality, transformed hierarchy score, and brokerage residual. Variables were standardized by column prior to clustering to place metrics measured on different scales on a comparable basis. Agglomerative hierarchical clustering was performed using Ward’s method. Missing values were not imputed; consequently, genes lacking a defined brokerage residual were excluded from the four-variable clustering analysis. Gene identity was used only as an observation label and did not contribute to clustering. HLDC assignments were not used as clustering variables and were therefore independent of dendrogram construction.

### Identification of Hierarchical Layers of Developmental Control

HLDC classification was interpretive rather than purely algorithmic. Network position was characterized using transformed hierarchy score and brokerage residual, whereas developmental function was assigned independently from published experimental studies of wing development. These sources of information were then integrated to assign genes to the five mechanistic HLDC categories defined below.

HLDC assignments were not defined by dendrogram membership or by the five quantile-based connectivity levels. Hierarchical clustering was instead used as an independent multivariate analysis to evaluate whether recurrent topological regimes corresponded to the biologically interpreted HLDC classes.

The five HLDC categories represent distinct mechanistic roles in the progressive transformation of developmental information into morphology:

### HLDC I: Organizer Centers

Genes involved in establishing global positional information through long-range developmental signaling.

### HLDC II: Signaling Scaffolds

Genes involved in interpreting and transmitting extracellular developmental signals through intracellular regulatory pathways.

### HLDC III: Interface Coordinators

Genes characterized by elevated brokerage capacity and developmental roles connecting partially autonomous developmental modules.

### HLDC IV: Pattern Implementers

Genes involved in specifying regional developmental identities and converting positional information into anatomical structures.

### HLDC V: Local Modulators

Genes involved in refining developmental outputs through localized regulation of signaling intensity, boundaries, and developmental precision.

Classification was performed using a combination of transformed connectivity hierarchy score, brokerage residual, and manual developmental annotation based on published functional studies.

### Developmental function annotation

Genes were manually annotated according to established developmental roles in wing development using primary literature and comprehensive reviews of Drosophila appendage patterning and signaling (Campbell and Tomlinson, 1998; Tanimoto *et al*., 2000; Irvine and Rauskolb, 2001; Barrio and de Celis, 2004; Schwank, Restrepo and Basler, 2008; Ruiz-Losada *et al*., 2018). Functional categories included:

⍰ Morphogen signaling
⍰ Growth regulation
⍰ Compartment and boundary specification
⍰ Selector gene activity
⍰ Patterning transcription factors
⍰ Pathway modulation
⍰ Tissue refinement and differentiation

Functional annotations were integrated with network metrics to determine whether connectivity position corresponded to distinct developmental mechanisms.

### Statistical analyses

Associations between degree and betweenness centrality were evaluated using linear regression of log-transformed values, with regression residuals used to quantify brokerage. Stability of network metrics across STRING confidence thresholds was evaluated using Spearman rank correlations calculated among genes retained in each pairwise comparison (Hagberg, Swart and Schult, 2008; Virtanen *et al*., 2020). Multivariate structure among network metrics was examined using principal component analysis and hierarchical clustering. Hierarchical clustering was performed on column-standardized degree centrality, betweenness centrality, transformed hierarchy score, and brokerage residual using Ward’s method. Multivariate analyses were performed in JMP Pro 19 (Jones and Sall, 2011). Genes lacking estimable brokerage residuals were excluded from analyses requiring this metric but retained in analyses of degree, betweenness, and connectivity hierarchy.

### AI tool usage

OpenAI ChatGPT (GPT-5 series) was used during this study to assist with the development and refinement of Python code for network analysis and data visualization, and organization of analytical outputs, and manuscript preparation. During manuscript preparation, the tool was used to assist with restructuring and editing author-generated scientific content for clarity and consistency. All analytical approaches, biological interpretations, developmental classifications and conclusions were determined by the author, and all AI-assisted code, analyses, and manuscript text were reviewed and verified by the author. No generative AI tools were used to generate or alter primary research data.

## Acknowledgements

The authors would like to thank Fred Nijhout and Richard Gawne for helpful feedback throughout the genesis of this project. We would also like to thank High Point University for supporting summer research students through SuRPS.

## Data and Resource Availability

All relevant data and details of resources can be found within the article and its supplemental information.

---

**Fig. S1.**
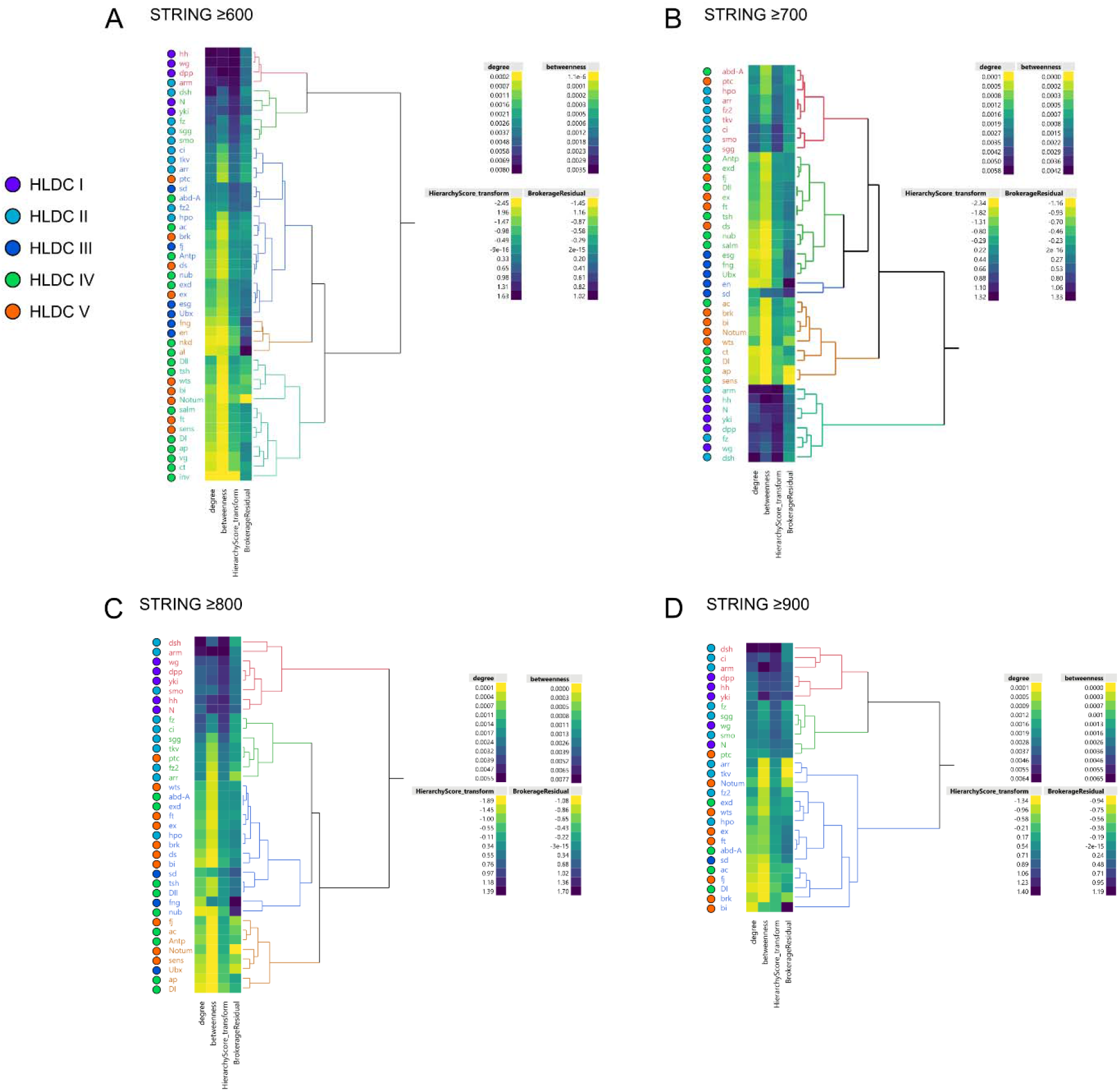
Hierarchical organization of wing developmental genes across STRING interaction-confidence thresholds. Hierarchical clustering of wing developmental genes was performed independently using STRING combined-score thresholds of (A) 600, (B) 700, (C) 800 and (D) 900. Clustering was based on column-standardized degree centrality, betweenness centrality, transformed hierarchy score and brokerage residual using Ward’s method. Colored circles indicate independently assigned Hierarchical Layers of Developmental Control (HLDCs) and were not used to construct the dendrograms. The broad multivariate organization of the network was retained across interaction-confidence thresholds, including separation of highly connected genes from lower-connectivity regions, although individual branch relationships and cluster membership varied with threshold. This variation is consistent with the greater threshold sensitivity of shortest-path and brokerage relationships relative to the overall connectivity hierarchy (Table S1). Genes lacking a defined brokerage residual because of zero betweenness were excluded from the corresponding clustering analysis.

**Fig. S2.**
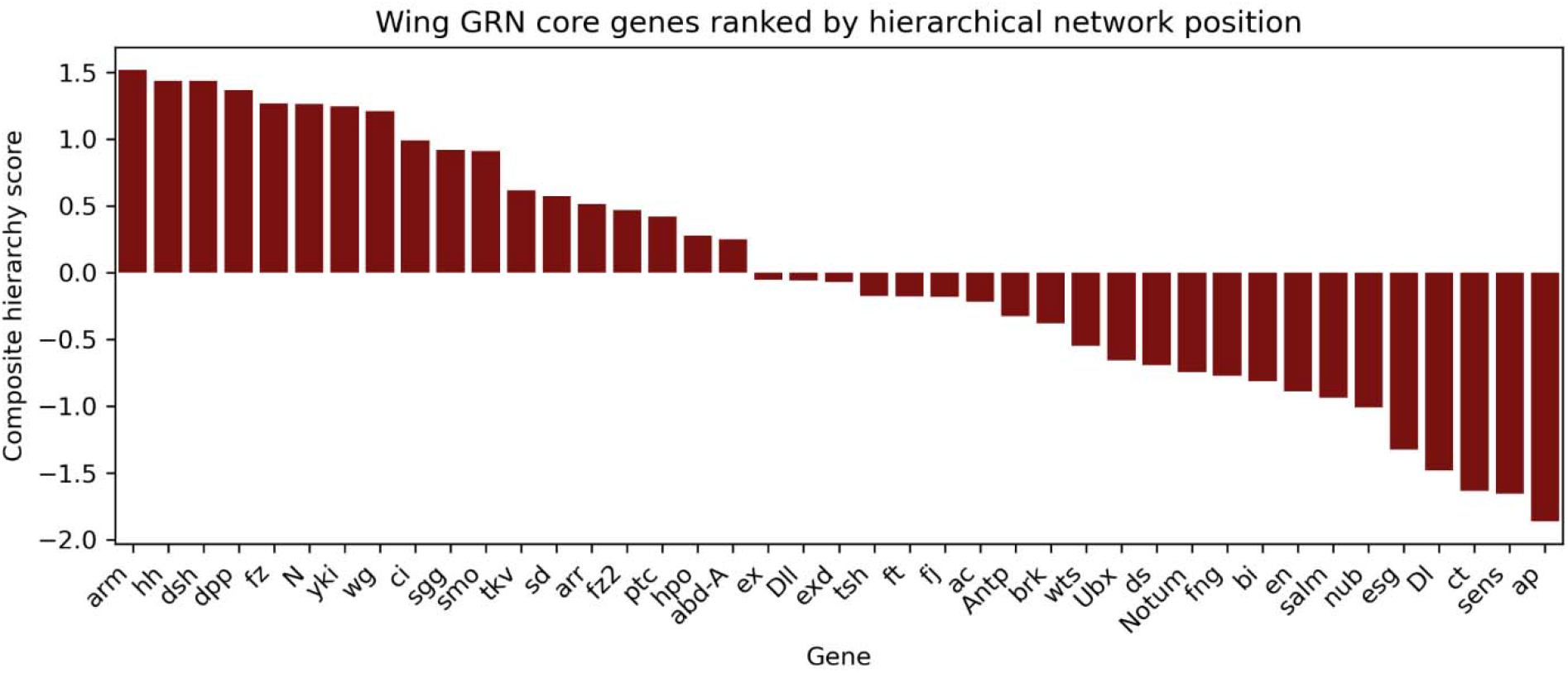
Composite hierarchy scores reveal a continuous core-to-periphery organization among wing developmental genes. Genes are ranked by transformed hierarchy score, calculated as the mean of standardized log-transformed degree and betweenness centrality. Positive values identify genes with greater combined connectivity and shortest-path centrality relative to the analyzed gene set, whereas negative values identify progressively more peripheral network positions. The continuous distribution illustrates the broad connectivity hierarchy underlying the network and was used to characterize network position independently of HLDC classification.

**Fig. S3.**
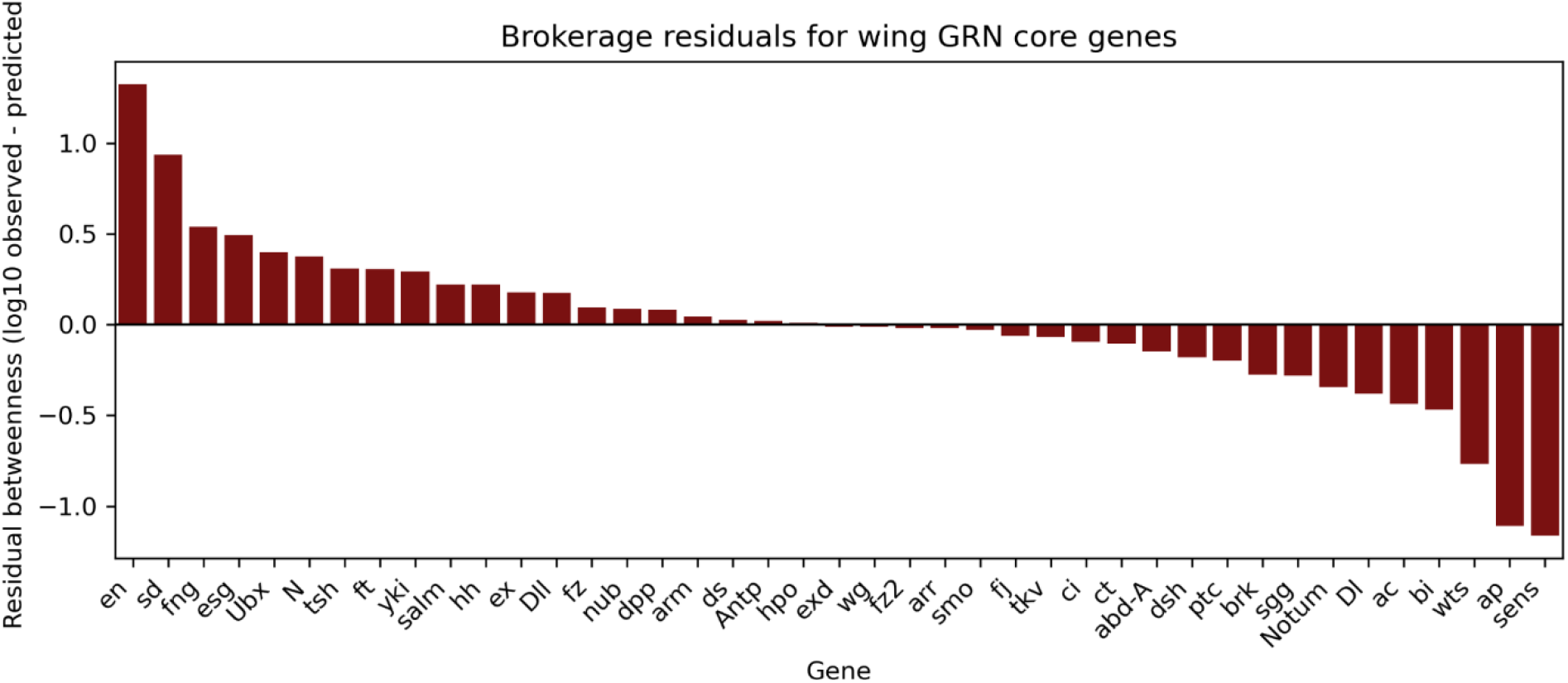
Brokerage residuals identify wing developmental genes with disproportionate betweenness relative to their connectivity. Genes are ranked by brokerage residual, calculated as the difference between observed log-transformed betweenness centrality and that predicted from log-transformed degree. Positive residuals identify genes occupying more shortest paths than expected from their overall connectivity, whereas negative residuals indicate lower betweenness than expected. High positive residuals for genes including *en*, *sd*, *fng*, *esg* and *Ubx* reveal a topological dimension distinct from the composite connectivity hierarchy and identify candidate bridging positions within the developmental network. Genes for which brokerage residuals could not be calculated because of zero betweenness are not shown.

**Table S1.** Robustness of wing developmental network topology across STRING interaction-confidence thresholds. Spearman rank correlations compare the relative network positions of wing developmental genes across STRING combined-score thresholds of 600, 700, 800, and 900. Correlations were calculated using genes present at both thresholds in each pairwise comparison. Brokerage residual correlations used pairwise-complete observations because residual brokerage was undefined for genes with zero betweenness. n, number of genes included in each comparison; ρ, Spearman rank correlation coefficient.

| Table S1. Robustness of wing developmental network |  |  |  |
| --- | --- | --- | --- |
| **A. Transformed hierarchy score** |  |  |  |
| Threshold comparison | n | Spearman $\rho$ | P |
| 600 vs 700 | 45 | 0.943 | 3.37E-22 |
| 600 vs 800 | 44 | 0.880 | 3.39E-15 |
| 600 vs 900 | 39 | 0.860 | 2.35E-12 |
| 700 vs 800 | 44 | 0.943 | 9.68E-22 |
| 700 vs 900 | 39 | 0.924 | 5.63E-17 |
| 800 vs 900 | 39 | 0.899 | 7.34E-15 |
| **B. Degree centrality** |  |  |  |
| Threshold comparison | n | Spearman $\rho$ | P |
| 600 vs 700 | 45 | 0.967 | 2.75E-27 |
| 600 vs 800 | 44 | 0.919 | 1.52E-18 |
| 600 vs 900 | 39 | 0.895 | 1.67E-14 |
| 700 vs 800 | 44 | 0.969 | 2.93E-27 |
| 700 vs 900 | 39 | 0.940 | 6.52E-19 |
| 800 vs 900 | 39 | 0.951 | 2.22E-20 |
| **C. Betweenness centrality** |  |  |  |
| Threshold comparison | n | Spearman $\rho$ | P |
| 600 vs 700 | 45 | 0.890 | 2.91E-16 |
| 600 vs 800 | 44 | 0.788 | 2.23E-10 |
| 600 vs 900 | 39 | 0.754 | 3.08E-08 |
| 700 vs 800 | 44 | 0.887 | 1.02E-15 |
| 700 vs 900 | 39 | 0.813 | 3.31E-10 |
| 800 vs 900 | 39 | 0.763 | 1.62E-08 |
| **D. Brokerage residual** |  |  |  |
| Threshold comparison | n | Spearman $\rho$ | P |
| 600 vs 700 | 41 | 0.578 | 7.70E-05 |
| 600 vs 800 | 37 | 0.282 | 9.07E-02 |
| 600 vs 900 | 28 | 0.442 | 1.86E-02 |
| 700 vs 800 | 37 | 0.541 | 5.45E-04 |
| 700 vs 900 | 28 | 0.415 | 2.81E-02 |
| 800 vs 900 | 28 | 0.697 | 3.73E-05 |

